# Pre-Steady-State Kinetic Characterization of an Antibiotic-Resistant Mutation of *Staphylococcus aureus* DNA Polymerase PolC

**DOI:** 10.1101/2022.10.04.510889

**Authors:** Rachel Nelson-Rigg, Sean P. Fagan, William J. Jaremko, Janice D. Pata

## Abstract

The emergence and spread of antibiotic resistance in bacterial pathogens are serious and ongoing threats to public health. Since chromosome replication is essential to cell growth and pathogenesis, the essential DNA polymerases in bacteria have long been targets of antimicrobial development, although none have yet advanced to the market. Here we use transient-state kinetic methods to characterize the inhibition of the PolC replicative DNA polymerase from *Staphylococcus aureus* by ME-EMAU, a member of the 6-anilinouracil compounds that specifically target PolC enzymes, which are found in low-GC content Gram-positive bacteria. We find that ME-EMAU binds to *S. aureus* PolC with a dissociation constant of 14 nM, more than 200-fold tighter than the previously reported inhibition constant, which was determined using steady-state kinetic methods. This tight binding is driven by a very slow off rate, 0.006 s^-1^. We also characterized the kinetics of nucleotide incorporation by PolC containing a mutation of phenylalanine 1261 to leucine (F1261L). The F1261L mutation decreases ME-EMAU binding affinity by at least 3500-fold, but also decreases the maximal rate of nucleotide incorporation by 11.5-fold. This suggests that bacteria acquiring this mutation would be likely to replicate slowly and be unable to out-compete wild-type strains in the absence of inhibitor, reducing the likelihood of the resistant bacteria propagating and spreading resistance.

## INTRODUCTION

The World Health Organization has placed antibiotic-resistant strains of *Staphylococcus aureus* on the list of priority pathogens for the research and development of new antibiotics (1). Antibiotics that directly target the replicative DNA polymerases, those that are primarily responsible for duplicating the bacterial chromosome, represent a novel therapeutic strategy. DNA polymerase III (Pol III) replicates the bulk of the bacterial genome and is essential for cell survival and pathogenesis (2). PolC is the catalytic subunit of Pol III in Gram-positive bacteria with low GC-content, while DnaE serves that role in Gram-negative bacteria (3, 4). Both enzymes are members the C-family of DNA polymerases (5). C-family DNA polymerases are only found in bacteria, and structural studies (6–8) have shown that they are not related to any of the eukaryotic replicative DNA polymerases, reducing the chances that inhibitors targeting the C-family polymerases would have off-target effects in humans.

Specific inhibitors of PolC were first identified in the 1960’s and are still being actively developed for the treatment of Gram-positive bacterial infections (9, 10). In fact, a phase 2 clinical trial of the PolC inhibitor ACX-362E (Ibezapolstat) is in progress for treatment of *Clostridium difficile* (11). These compounds compete with deoxynucleotide triphosphate (dNTP) binding at the active site of PolC by forming hydrogen bonds with the nucleotide in the templating strand of DNA that is the next to be copied by the polymerase. The 3’-ethyl-4’methylaniliouracils (EMAUs), an earlier class of compounds that work in the same way, are potent inhibitors of PolC and comprise two functional domains (Fig. 1A): the uracil domain forms hydrogen bonds that mimic the pairing of dGTP with dC while the aryl domain interacts with the polymerase (12). The 2-methoxyethyl derivative of EMAU (ME-EMAU) is bactericidal at a concentration of four-times the minimal inhibitory concentration of 16 µg/mL for the strain of *S. aureus* tested (13). At a low frequency, resistance to EMAU arises due to mutation of a single amino acid substitution, F1261L in *S. aureus* PolC (14).

**Figure 1.**
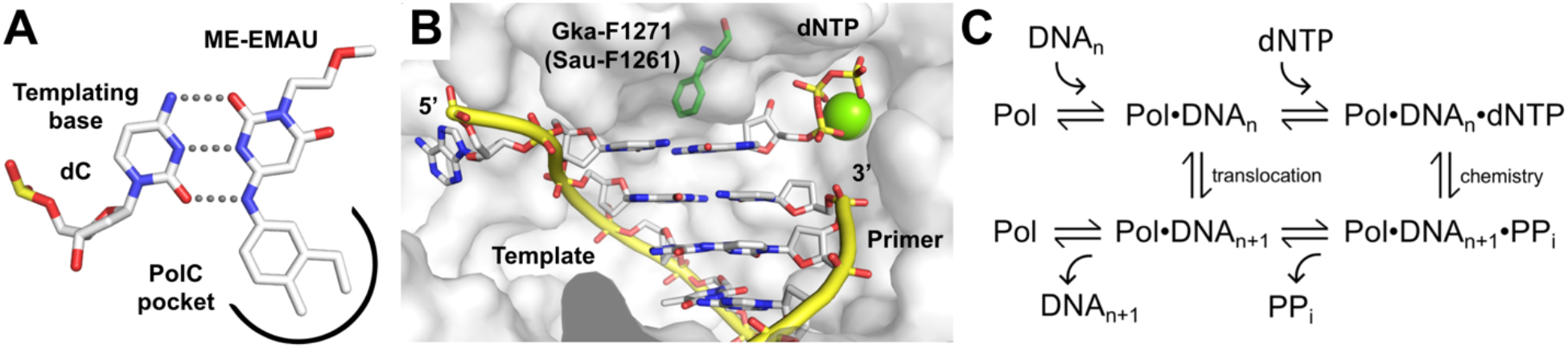
Mechanisms of ME-EMAU inhibition, resistance and DNA synthesis. (A) ME-EMAU and other anilinouracil derivatives compete with dGTP binding to cytosine in the template DNA by forming hydrogen bonds. This mimics a Watson-Crick basepair and sequesters the polymerase in an unproductive complex. (B) Active site of *G. kaustophilus* (Gka) PolC (PDB code 3F2B) showing the location of primer-template DNA and dNTP relative to residue F1271, which is equivalent to *S. aureus* (Sau) PolC residue F1261. (C) Minimal kinetic reaction pathway for DNA polymerases. Pol (DNA Polymerase); DNA_n_ (unextended DNA); dNTP (deoxynucleotide triphosphate); DNA_n+1_ (DNA extended by one nucleotide); PPi (pyrophosphate).

During catalysis, enzymes adopt multiple states along a reaction pathway and each structural state represents a different drug target (15). To effectively target the different reaction intermediates, a comprehensive kinetic description of these states is essential and can be achieved using transient-state rather than steady-state kinetic measurements. We have recently determined the mechanism *S. aureus* PolC using transient-state kinetics (16, 17), the first detailed kinetic characterization of any C-family DNA polymerase. The structure of a thermophilic homolog of *S. aureus* PolC in a ternary complex that is poised for catalysis (Fig. 1B) provides detailed information about the contacts between the polymerase and its substrates, the primer-template DNA and the incoming dNTP (8). DNA polymerases share a common two-metal-ion mechanism for phosphodiester bond formation (18), with the different activities of polymerases arising primarily because of differences in substrate binding affinity and the rates of each step in the reaction pathway (Fig. 1C).

Here we use transient-state kinetics to characterize the interaction of ME-EMAU with *S. aureus* PolC and the properties of the F1261L PolC mutation that confers resistance to ME-EMAU. We find that ME-EMAU binds to PolC much more tightly than predicted based on prior steady-state kinetic measurement and that the F1261L mutation significantly impairs the polymerase activity of PolC, thus reducing the fitness of the enzyme.

## RESULTS

PolC has two catalytic domains: the polymerase domain that synthesizes DNA and a 3’-to-5’ exonuclease proofreading domain that increases the fidelity of DNA replication by removing mis-incorporated nucleotides. The exonuclease domain of both the wild-type and mutant enzymes was inactivated by four aspartate-to-alanine substitutions so that we could characterize the polymerase activity independently from any DNA degradation (17). For simplicity, we refer to these enzymes as PolC-WT and PolC-F1261L, even though they both contain the exonuclease-inactivating mutations. In all assays, we have also included the *S. aureus* β-clamp (DnaN), a key component of the bacterial replisome that acts as a processivity factor by encircling the DNA duplex and binding to the C-terminal tail of PolC, which topologically tethers the polymerase to the circular chromosomal DNA. Even on a short, linear DNA substrate, the β-clamp increases the affinity of PolC for DNA by four-fold (17). The DNA substrate used in this study (Fig. 2A) has a duplex length of 29 basepairs, a length that is sufficient to allow binding of both PolC and the β-clamp (8, 19), and a single-stranded template region of 39 nucleotides (Fig. 2A). This substrate differs from the one that we used in our prior study of PolC (17) by a single nucleotide; the FAM-labeled primer strand is one nucleotide shorter, such that the first templating base is a cytosine that will pair either with dGTP as the next-correct nucleotide or with the ME-EMAU inhibitor.

**Figure 2.**
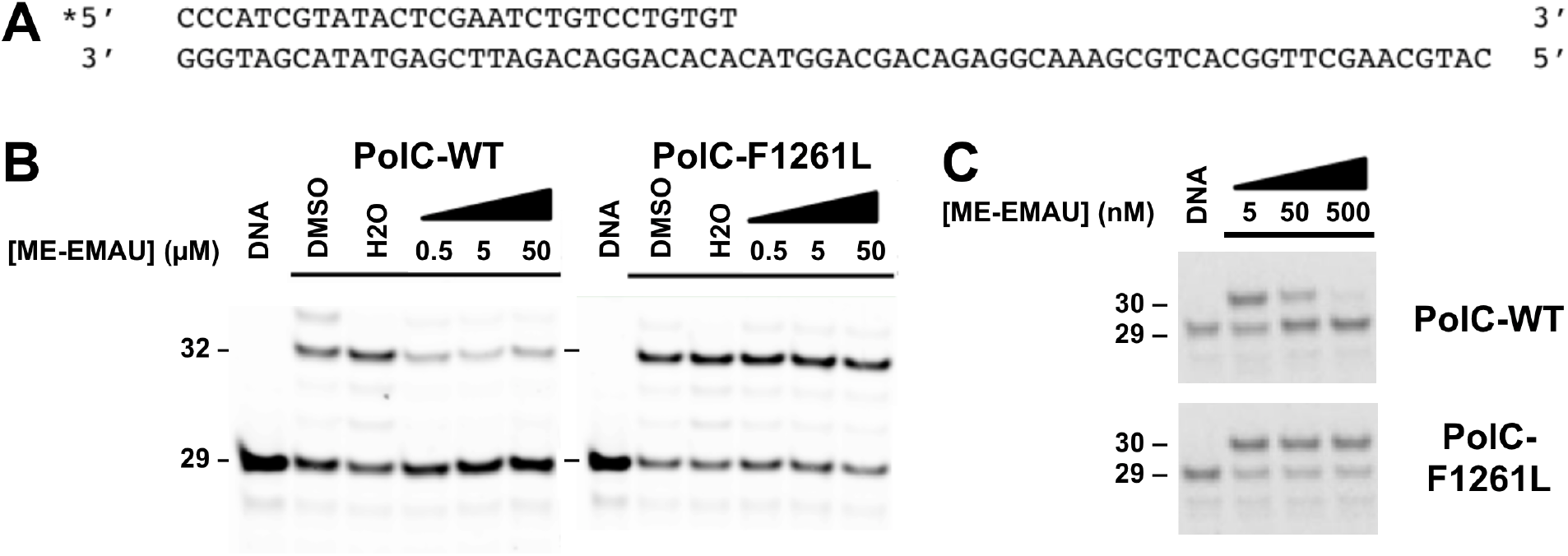
The F1261L mutation in *S. aureus* PolC confers resistance to ME-EMAU. (A) The DNA substrate used in this study. The primer strand is labeled at the 5’ end with 6-carboxyfluorescein (FAM; *). (B) and (C) Primer extension reactions performed in the presence of increasing concentrations of ME-EMAU. Reactions in (B) contained 0.5, 5 or 50 µM ME-EMAU, 10 µM each nucleotde (dGTP, dTTP and dATP)(and were incubated for 60 seconds. Reactions in (C) contained 5, 50 or 500 nM ME-EMAU, 10 µM dGTP and were incubated for 1 second. All reactions contained 500 nM each PolC and DNA. Control reactions contained either DMSO or water (H2O) in place of the inhibitor. Lanes with DNA alone show the unextended primer.

### ME-EMAU binds tightly to PolC•DNA and dissociates slowly

Initial experiments showed that PolC-WT is more sensitive to ME-EMAU than we had anticipated based on the previously reported inhibition constant (K_i_) of ∼3 µM (13). Varying concentrations of ME-EMAU (0.5, 5, 50 µM) were preincubated with PolC and DNA, to allow formation of the inhibited PolC•DNA•ME-EMAU complex. The first three correct nucleotides (dGTP, dTTP and dATP) were then added to initiate DNA synthesis and the reaction was allowed to proceed for 60 seconds. DNA synthesis by PolC-WT was substantially reduced in the presence of ME-EMAU, but all three concentrations tested showed comparable amounts of inhibition (Fig. 2B, left panel). In subsequent assays that used lower concentrations of ME-EMAU (5, 50 and 500 nM), a shorter reaction time (1 second) and a single nucleotide (dGTP), we observed concentration-dependent inhibition (Fig. 2C, top panel), suggesting that the K_i_ in our assays was in the low nanomolar range.

To determine the affinity of ME-EMAU for the PolC•DNA binary complex accurately, we turned to transient-state kinetics in which the concentrations of enzyme and substrate are comparable, thus facilitating detection of reaction products from a single enzyme turnover (20, 21). PolC displays biphasic reaction kinetics, with an initial burst of rapid synthesis occurring in the first 10-20 ms of the reaction that is followed by a slower linear phase (16, 17). The amplitude of the burst phase corresponds to the amount of active, pre-assembled PolC•DNA complex that is capable of DNA synthesis when the reaction is initiated with dGTP. The slower linear phase after the burst results from subsequent catalytic cycles and indicates the presence of a slow step after catalysis that limits the rate of multiple enzyme turnovers.

The binding dissociation constant (K_D_) for the primer-template DNA and dNTP substrates and the K_i_ for inhibitor were determined by varying the concentration of each reaction component independently of the others and measuring a time course that covers both phases of the reaction. PolC and DNA were incubated together, in the presence or absence of ME-EMAU, prior to addition of dGTP (Fig. 3A). In all reactions, the total concentration of PolC was 500 nM and the incubation time was varied from 2 to 80 ms. The amplitude of the fast phase of the reaction increased with increasing DNA concentration (50-1600 nM; Fig. 3B) and decreased with increasing ME-EMAU concentration (2.5-200 nM; Fig. 3C). Both the amplitude and rate of the fast phase of the reaction increased with increasing dGTP concentration (0.5-75 µM; Fig. 3D). To measure the apparent rate of ME-EMAU binding, the reaction scheme was altered (Fig. 3E) such that PolC and DNA were pre-incubated together, then a saturating concentration of ME-EMAU (25 µM) was added and allowed to interact with the PolC•DNA complex for varying times (2-1000 ms) prior to the addition of a saturating concentration of dGTP (250 µM) to initiate the reaction, which was allowed to proceed for 28 ms before quenching. The amount of DNA synthesis decreased as the incubation time with ME-EMAU increased (Fig. 3F). The results from these experiments were fit globally to the reaction pathway shown (Fig. 3G) using KinTek Explorer (22, 23) to determine a single set of kinetic constants that best fit all the data (Table 1).

**Table 1.** Pre-steady-state kinetic parameters for wild-type and F1261L Sau-PolC.

| **Globally Fit Parameters** | **WT** | **F1261L** | **WT/F1261L** |
| --- | --- | --- | --- |
|  | <u>Best Fit ± SE</u><br><u>(95% CI)</u> | <u>Best Fit ± SE</u><br><u>(95% CI)</u> | <u>Fold Change</u> |
| ***DNA binding:*** |  |  |  |
| $k_{+1}$ ($\mu\text{M}^{-1}\text{s}^{-1}$) | $6.2 \pm 0.1$<br>(5.8 – 6.6) | $3.7 \pm 0.4$<br>(2.7 – 5.8) | 1.7 |
| $k_{-1}$ ($\text{s}^{-1}$) | 1.1 (fixed) | $0.42 \pm 0.01$<br>(0.39 – 0.45) | 2.6 |
| ***Nucleotide binding:*** |  |  |  |
| $k_{+2}$ ($\mu\text{M}^{-1}\text{s}^{-1}$) | 100 (fixed) | 100 (fixed) | – |
| $k_{-2}$ ($\text{s}^{-1}$) | $1440 \pm 87$<br>(1160 – 1860) | $274 \pm 104$<br>(0 – 707) | 5.2 |
| ***Polymerization rate:*** |  |  |  |
| $k_{+3}$ ($\text{s}^{-1}$) | $1196 \pm 35$<br>(1070 – 1360) | $104 \pm 17$<br>(60 – 168) | 11.5 |
| $k_{-3}$ ($\text{s}^{-1}$) | $122 \pm 11$<br>(75 – 166) | $51 \pm 18$<br>(0.05 – 124) | 2.4 |
| ***ME-EMAU binding:*** |  |  |  |
| $k_{+5}$ ($\mu\text{M}^{-1}\text{s}^{-1}$) | $0.42 \pm 0.02$<br>(0.35 – 0.47) | nd | – |
| $k_{-5}$ ($\text{s}^{-1}$) | $0.006 \pm 0.001$<br>(0.005 – 0.007) | nd | – |
| **Calculated Parameters** | **WT** | **F1261L** | **WT/F1261L** |
|  | <u>Best Fit ± Std Error</u> | <u>Best Fit ± Std Error</u> | <u>Fold Change</u> |
| $K_D^{\text{DNA}}$ ($\mu\text{M}$) | $0.18 \pm 0.02$ | $0.11 \pm 0.01$ | 1.6 |
| $K_D^{\text{dGTP}}$ ($\mu\text{M}$) | $14.4 \pm 0.9$ | $2.7 \pm 1.0$ | 5.2 |
| $k_{\text{pol}}$ ($\text{s}^{-1}$) | $1318 \pm 37$ | $155 \pm 106$ | 8.5 |
| $k_{\text{pol}}/K_D^{\text{dGTP}}$ ($\mu\text{M}^{-1}\text{s}^{-1}$) | $92 \pm 6$ | $57 \pm 44$ | 1.6 |
| $K_D^{\text{ME-EMAU}}$ ($\mu\text{M}$) | $0.014 \pm 0.003$ | – | – |

**Figure 3.**
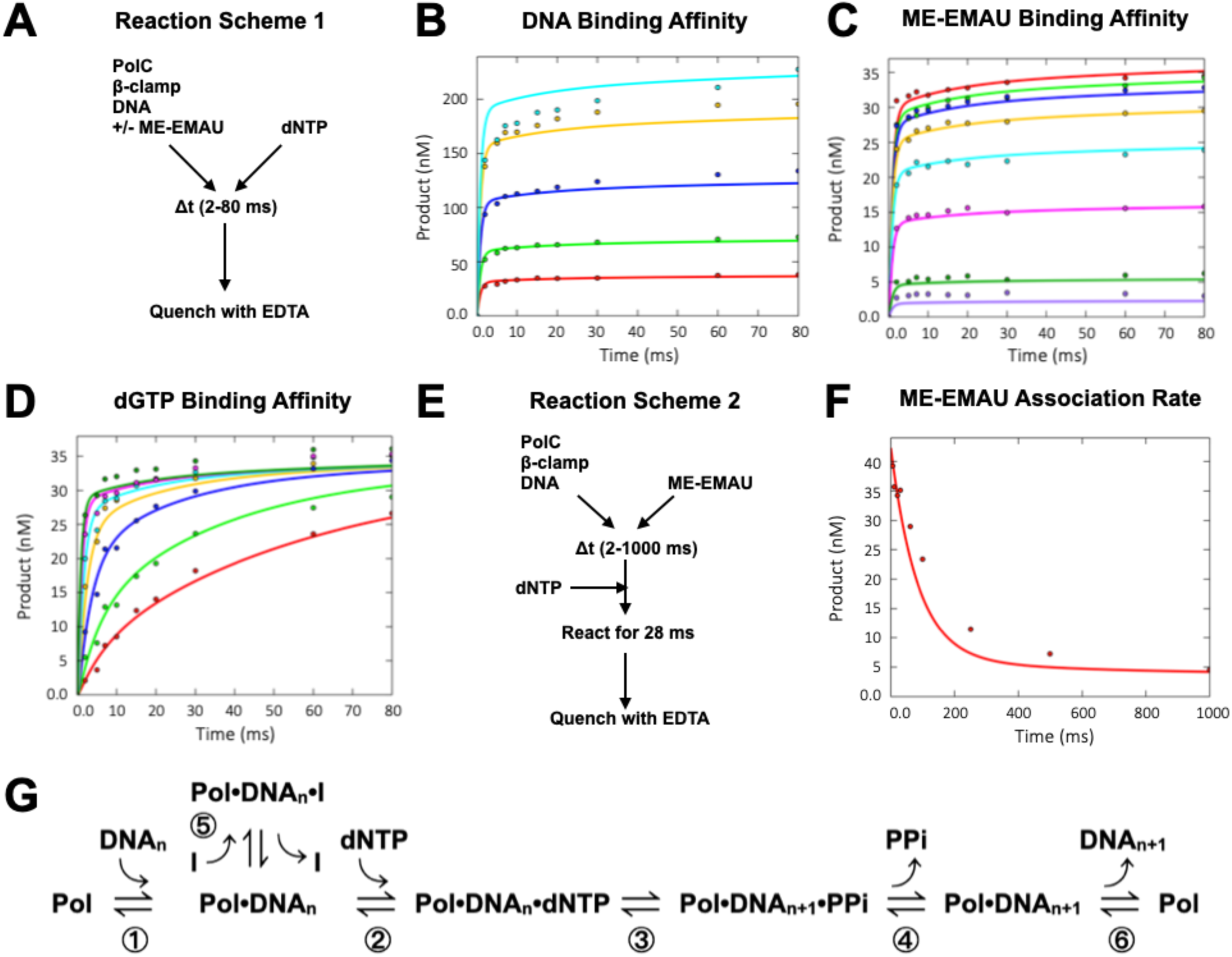
Single-nucleotide incorporation kinetics of PolC-WT in the absence and presence of ME-EMAU. (A) Reaction scheme used in (B-D) to determine (B) DNA binding affinity, (C) ME-EMAU binding affinity, and (D) nucleotide binding affinity. Polymerase (0.5 µM) was pre-incubated with DNA (50-1600 nM in B; 50 nM in C and D) in the absence (B and C) or presence (D) of ME-EMAU (2.5-200 nM) prior to mixing with dGTP (100 µM in B and D; 0.5-75 µM in C) for varying times (2-80 ms). (E) Reaction scheme used in (F) to determine the rate of ME-EMAU binding to the PolC•DNA complex. Polymerase (0.5 µM) was pre-incubated with DNA (50 nM), mixed with ME-EMAU (25 µM) for varying times (2-1000 ms) prior to the addition of a saturating concentration of dGTP (125 µM) for 28 ms. All concentrations given are the final total concentrations in the reaction after mixing. Curves in (B-D and F) were derived by globally fitting the data to the reaction pathway shown in (G) using KinTek Explorer. Kinetic constants derived from the global fitting are shown in Table 1.

These experiments demonstrate that ME-EMAU binds to PolC-WT with a K_D_^EMAU^ of 14 nM,

>200-fold more tightly than determined from steady-state measurements (13), and dissociates very slowly, with an off rate (k_off_^EMAU^ = k_-6_) of 0.006 s^-1^. The substrates bind with a K_D_^DNA^ of 180 nM and a K_D_^dGTP^ of 14.4 µM, with ∼49% of the enzyme capable of forming an active PolC•DNA complex that can catalyze product formation upon dGTP addition. Additionally, release of pyrophosphate (PPi) is rate-limiting, with a k_off_^PPi^ (k_4_) of 66.5 s^-1^ calculated from the global fitting, which is ∼20-fold slower than the maximal polymerization rate (k_pol_) of 1318 s^-1^, the sum of the forward and reverse reaction rates. All these parameters are comparable to those that we determined previously for the single-nucleotide incorporation of dTTP and dATP (17).

### ME-EMAU-resistant mutant PolC-F1261L incorporates dNTP slowly

PolC-F1261L was resistant to ME-EMAU, even at a concentration of 50 µM (Fig. 2B, right panel; Fig. 2C, bottom panel), indicating that the single mutation decreases inhibitor binding affinity by at least 3500-fold. To determine if the F1261L mutation altered the kinetic parameters of PolC, we used the same transient-state methods that we used for PolC-WT. In addition to measuring the binding affinities for DNA and dGTP (Figs. 4A and 4B), we determined the rate of DNA dissociation from PolC-F1261L using a modified reaction scheme (Fig. 4C) in which a pre-incubated mixture of enzyme and labeled DNA was mixed with a 500-fold excess of unlabeled trap DNA for varying times (2 ms to 120 sec) prior to the addition of dGTP. Upon dGTP addition, only the enzyme still bound to the labeled DNA would produce visible reaction products, resulting in a decreased reaction amplitude as the mixing time increased (Fig. 4D).

**Figure 4.**
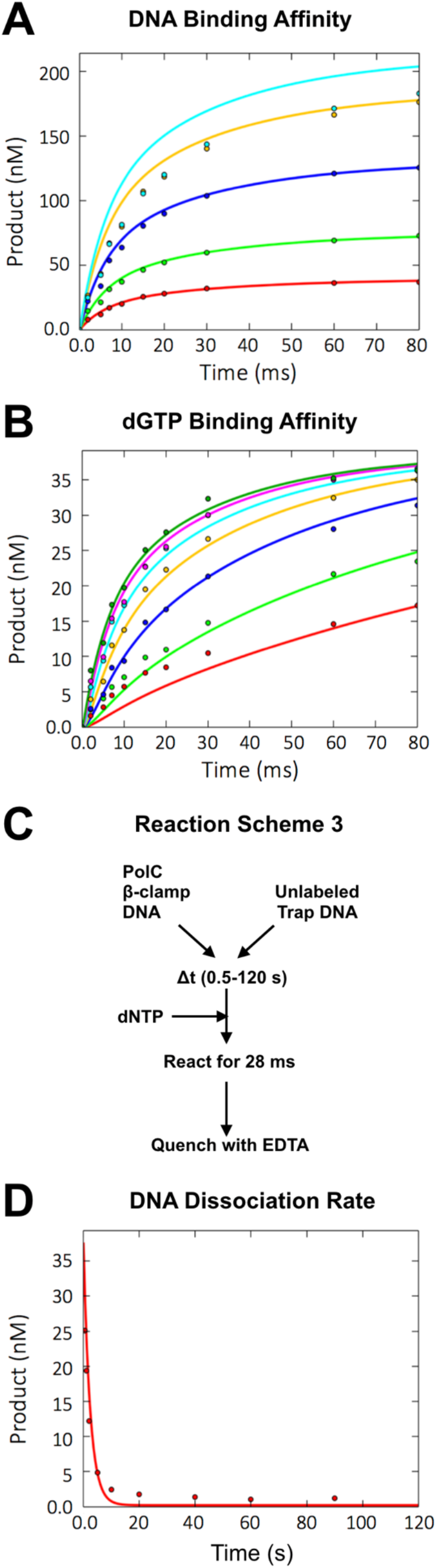
Single-nucleotide incorporation kinetics of PolC-F1261L. The affinities of PolC-F1261L for (A) DNA and (B) dGTP were determined as for the wild-type enzyme (Fig. 3A-C). (C) Reaction scheme used in (D) to determine the rate of DNA dissociation from PolC-F1261L. PolC (0.5 µM) was pre-incubated with labeled DNA (50 nM), mixed first with unlabeled DNA (25 µM) for varying times (0.08-9 s) prior to the addition of dGTP (1 mM) for 28 ms. All concentrations given are the final total concentrations in the reaction after mixing. Curves in A, B and D were derived by globally fitting the data to the reaction pathway shown in Fig. 3G using KinTek Explorer. Kinetic constants derived from the global fitting are shown in Table 1.

When the data for PolC-F1261L were fit globally, we found two substantial differences between the mutant and wild-type polymerases (Table 1). The mutant enzyme has a maximal polymerization rate that is 8.5-fold slower than wild-type (k_pol_ 155 s^-1^ vs. 1318 s^-1^) but binds 5.2-fold more tightly to dGTP (K_D_^dGTP^ 2.7 µM vs. 14.4 µM). Taken together, these parameters result in a 1.6-fold decrease in nucleotide incorporation efficiency (k_pol_/ K_D_^dGTP^) by the mutant enzyme. Comparable to the wild-type enzyme, ∼43% of PolC-F1261L was capable of forming an active complex with DNA, indicating that the mutation does not have a significant effect on the overall activity of the enzyme.

## DISCUSSION

This transient-state kinetic analysis of *S. aureus* PolC demonstrates that ME-EMAU is a very potent inhibitor, binding 1000-fold more tightly to the PolC•DNA complex than does dGTP and dissociating 220,000-fold more slowly than the maximal rate of polymerization. Steady-state assays are dominated by the rate-limiting step(s) in a reaction cycle. In the case of single nucleotide incorporation by DNA polymerases, the rate-limiting step is typically the dissociation of the product DNA from the post-chemistry binary complex (Pol•DNA_n+1_). However, during processive DNA synthesis in vivo, the product DNA does not dissociate after each round of nucleotide incorporation. Instead, the polymerase translocates along the DNA to catalyze the addition of subsequent nucleotides. Thus, steady-state assays interrogating the kinetics of DNA polymerases typically inform us about a catalytic step that is not physiologically relevant. Moreover, such multiple-turnover assays do not typically provide information about the dNTP binding and incorporation steps that are often of the most interest for understanding the mechanism of nucleotide incorporation and replication fidelity. This effect is clearly illustrated in the 200-fold difference between the 14 nM dissociation constant for ME-EMAU that we report here and the 2.8 µM inhibition constant reported previously (13). In the case of PolC, multiple-turnover reactions are limited by rates of both DNA and PPi dissociation (17). In the previous steady-state assays of ME-EMAU, reaction times were 10 minutes, which we now know is 5-fold longer than the half-life of ME-EMAU binding to PolC. Even though the dissociation rate is extremely slow (0.006 s^-1^), the maximal rate of nucleotide incorporation (1322 s-1) is more than five orders of magnitude faster, causing the apparent inhibition of PolC to diminish over time, as is seen in the apparently weaker level of inhibition with a reaction time of 1 minute (Fig. 2B, left panel) vs. 1 second (Fig. 2C, top panel).

A key concern with any antibiotic is the potential for resistance mutations and the consequences of those mutations on the fitness of the resistant bacteria. The single amino acid substitution F1261L that confers ME-EMAU resistance on PolC decreases the maximal rate of nucleotide incorporation by 8.5-fold but the mutation also increases the affinity of the polymerase for nucleotides by 5.3-fold. These changes partially counteract each other such that the overall enzyme efficiency only decreases by 1.6-fold. However, the concentration of dNTPs in bacterial cells is 50-150 µM during exponential growth (24, 25), which is near saturation for both wild-type and mutant PolC. Under these conditions, the differences in nucleotide incorporation rate would have the most impact on bacterial fitness and we would expect cells containing the PolC-F1261L mutation to be at a substantial growth disadvantage.

The power of combining detailed structural and mechanistic kinetic information has been exemplified by the progression of antiretroviral therapies used to treat HIV-1 infection and AIDS (26). The first co-crystal structure of Nevirapine bound to HIV-1 reverse transcriptase (27, 28) and the kinetic mechanisms of nucleotide incorporation and inhibition by the enzyme (29, 30) provided the foundation for developing more effective generations of non-nucleoside reverse transcriptase inhibitors, including ones that are active against mutants that are resistant to Nevirapine. While transient-state kinetic methods would not replace steady-state assays for inhibitor screening, a detailed understanding of the kinetic pathway can help in the design of high-throughput screens that target individual reaction steps and can aid in optimizing initial hits in a screen, for example by comparing the binding and dissociation rates during the analysis of structure-activity relationships.

Compared to viral and bacteriophage replicative DNA polymerases, mechanistic studies of cellular replicative polymerases have lagged behind. Among the cellular replicative polymerases, the bacterial C-family polymerases have been particularly challenging to study, largely because of weak binding to DNA and the unique catalytic cycle of the enzyme, but the available structures of several essential C-family polymerases (7, 8, 31–36) and our transient-state kinetic characterization of *S. aureus* PolC (16, 17, 24) provide the framework needed for a mechanistic structure-function approach to developing antibiotics targeting bacterial DNA replication.

## MATERIALS AND METHODS

### Materials

DNA oligonucleotides were purchased from Integrated DNA Technologies Inc. (Coralville, IA, USA). Deoxynucleotides (dNTPs), ultrapure grade, and chromatography materials were purchased from GE Healthcare Biosciences (Schenectady, NY, USA).

### Plasmids

The *polC* gene from *S. aureus* strain COL was synthesized and codon-usage was optimized for expression in *Escherichia coli* by GenScript USA Inc. (Piscataway, NJ, USA) and cloned into pET28a(+) using restriction sites NcoI and BamHI, The resulting plasmid contained an N-terminal deca-histidine tag followed by a TEV cleavage site and contained the following mutations: F1261L in the polymerase domain, to confer resistance to ME-EMAU, together with D424A, E426A, D509A and D568A in the exonuclease domain, to prevent nucleic acid degradation. A plasmid containing the wild-type sequence at the active site was created using the Q5 site-directed mutagenesis kit from New England Biolabs (Ipswich, MA, USA), to reverse the resistance mutation at position 1261.

### Protein expression and purification

Plasmids containing the genes for wild-type and F1261L mutant PolC were transfected into Rosetta(DE3)pLysS *E. coli* and cells were grown in Terrific Broth with 100 μg/mL ampicillin, 17 μg/mL chloramphenicol, 1 mL/L glycerol, 660 mM sorbitol, and 2.5mM betaine at 37°C to an OD_600_ of 0.6 to 0.7. The temperature was then decreased from 37°C to 17°C and cells were induced with 0.5 mM IPTG and harvested 18 hours after induction. Protein was purified as described previously by chromatography over HisTrap HP charged with Nickel, Q-sepharose, heparin and Superdex 200 columns (16, 17). Purified protein was concentrated, flash-frozen in liquid nitrogen and stored at -80°C in 50 mM Tris-Cl pH 7.5, 250mM NaCl, 5mM EDTA, 10% glycerol, 5mM DTT. PolC concentration was determined by UV absorbance at 280 nm using a theoretical extinction coefficient of 113,000 M^-1^ cm^-1^. β-clamp was purified as described previously (17).

### DNA substrates

Oligonucleotide sequences for the polymerase substrate are shown in Fig. 2A. The primer strand, labeled at the 5’ end with 6-FAM, was annealed with a 1.1 molar excess of the unlabeled template strand in 10 mM Tris-HCl pH 8.0 and 50 mM NaCl by denaturing at 95°C and slowly cooling to room temperature.

### Polymerase assays

Polymerase reactions contained final concentrations of 25 mM MES-Tris (pH 7.5), 25 mM NaCl, 8 mM MgCl_2_, 2 mM DTT, 5% glycerol, 5 μM BSA, 0.5 µM total polymerase (PolC-WT or PolC-F161L), and 20 μM β-clamp monomer (10 µM dimer). All reactions were performed at room temperature and were terminated by the addition of an equal volume of quench buffer (250 mM EDTA, 50% formamide). Reactions with incubation times less than 1 minute were performed as quench-flow assays using a KinTek RQF-3 apparatus (KinTek Corp., Austin, TX, USA). Reaction products were separated on denaturing gels containing 7 M urea, 1xTBE, and 11 or 17% acrylamide (19:1) and run at 55°C. All gels were imaged on a Typhoon RGB (GE Life Sciences) scanner with an excitation wavelength of 488 nm (blue laser) and an emission cutoff of 525 nm. Products were quantitated by measuring the intensity of the band corresponding to primer extension relative to the amount of total labeled DNA (both unextended and extended) using ImageQuant software.

For multiple-nucleotide incorporation assays (Fig. 2B), PolC (WT or F1261L) was pre-incubated with DNA (500 nM final concentration each) and β-clamp in the presence of 0.05 to 50 µM ME-EMAU (dissolved in 100% DMSO) for 30 minutes. The final amount of DMSO in the reactions was kept constant at 10%. Reactions were initiated by the addition of dGTP, dTTP and dATP (the first three correct dNTPs for the substrate DNA) to a final concentration of 10 µM each and incubated for 60 s. For single-nucleotide incorporation assays (Figs. 2C, 3 and 4), the concentrations of primer-template DNA, dGTP, or ME-EMAU and the incubation time were varied depending on the experiment; final total concentrations during the extension reactions and reaction time are listed in the figure legends.

To determine the apparent rate of ME-EMAU binding to the PolC-WT•DNA complex (k_app_), a double-mixing experiment was performed (Fig. 3E and 3F) in which pre-incubated PolC-WT, DNA and β-clamp were mixed with a saturating concentration (25 µM final) of ME-EMAU for varying times (2 ms to 1 s) after which a saturating concentration (125 µM final) of dGTP was added and the reaction was allowed to proceed for 28 ms before quenching. A single exponential decay equation (Equation 1) was fit to the concentration of extended product (Y) to determine k_app_, where A_0_ is the amount of product formed in the absence of inhibitor, t is the mixing time, and c is a constant.

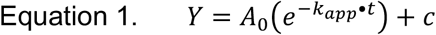

The rates of ME-EMAU binding and dissociation (k_on_ and k_off_) were calculated from the apparent rate of binding (k_app_) and the concentration of ME-EMAU using equations 2 and 3, where K_D,app_EMAU is the apparent dissociation constant for ME-EMAU, obtained from the experiment shown in Fig. 3C.

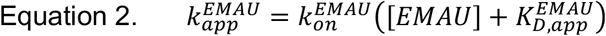

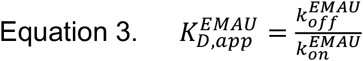

### Global analysis of kinetic data

Global fitting of the data by simulation was done using KinTek Explorer software version 10.0.200514 (22). The forward rate of dNTP binding was locked in at a diffusion limited rate of 100 μM s^-1^ and the reverse rate for PPi release was locked at a slow rate of 0.001 s^-1^, since rebinding of PPi to PolCWT is negligible under the reaction conditions used. The dissociation rate of PolC-WT from DNA was fixed at 1.1 s^-1^ based on previous kinetic characterization of the polymerase from a similar template where the primer strand was a single nucleotide longer. For PolC-F1261L, forward and reverse rates for PPi release were fixed at 62.7 s^-1^ and 0.01 s-1, respectively, according to our previous data demonstrating that PPi release is rate-limiting (17). One- and two-dimensional confidence contour analyses were performed using the Fitspace function (23).

## ACKLOWLEDGEMENTS

We thank Indrajit Lahiri for helpful discussions and critical reading of the manuscript, Tori Smiraglia for technical assistance, and Ken Johnson for generously sharing his expertise and insights into polymerase kinetics. We thank the following Wadsworth Center core facilities for providing expertise and material support: media and tissue culture, applied genomic technologies core, biochemistry, and protein expression.

